# Molecular Contribution to Embryonic Aneuploidy and Genotypic Complexity During Initial Cleavage Divisions of Mammalian Development

**DOI:** 10.1101/2020.07.24.220475

**Authors:** Kelsey E. Brooks, Brittany L. Daughtry, Brett Davis, Melissa Y. Yan, Suzanne S. Fei, Lucia Carbone, Shawn L. Chavez

## Abstract

Embryonic aneuploidy is highly complex, often leading to developmental arrest, implantation failure, or spontaneous miscarriage in both natural and assisted reproduction. Despite our knowledge of mitotic mis-segregation in somatic cells, the molecular pathways regulating chromosome fidelity during the error-prone cleavage-stage of mammalian embryogenesis remain largely undefined. Using bovine embryos and live-cell fluorescent imaging, we observed frequent micro-/multi-nucleation of anaphase lagging or mis-segregated chromosomes in initial mitotic divisions that underwent unilateral inheritance, re-fused with the primary nucleus, or formed a chromatin bridge with neighboring cells. A correlation between a lack of maternal and paternal pronuclei fusion (syngamy), multipolar cytokinesis, and uniparental genome segregation was also revealed and single-cell DNA-seq showed propagation of primarily non-reciprocal mitotic errors in embryonic blastomeres. Depletion of the mitotic checkpoint protein, BUB1B/BUBR1, resulted in micro-/multi-nuclei formation, atypical cytokinesis, chaotic aneuploidy, and disruption of the kinase-substrate network regulating mitotic progression and exit, culminating in embryo arrest prior to genome activation. This demonstrates that embryonic micronuclei sustain multiple fates, provides a mechanism for blastomeres with uniparental origins, and substantiates the contribution of defective checkpoint signaling and/or the inheritance of other maternally-derived factors to the high genotypic complexity afflicting preimplantation development in higher-order mammals.

## INTRODUCTION

Multiple studies across higher-order mammalian species, including humans, have established that *in vitro*-derived embryos suffer from remarkably frequent whole chromosomal losses and gains termed aneuploidy (Vanneste et al. 2009; Daughtry et al. 2019). Depending on the type and severity of the chromosome segregation error, many aneuploid embryos will undergo developmental arrest and/or result in early pregnancy loss if transferred. Estimates of embryonic aneuploidy *in vivo* are difficult to ascertain, but ~50-70% of spontaneous miscarriages following natural conception in women are diagnosed as karyotypically abnormal (Hassold et al. 1980; Schaeffer et al. 2004). Aneuploidy can arise either meiotically during gametogenesis, or post-zygotically from the mitotic cleavage divisions of preimplantation development. Although significant effort has been put forth to identify specific contributors to meiotic chromosome mis-segregation, particularly with advanced maternal age (Webster and Schuh 2017; Schneider and Ellenberg 2019), much less is known about the molecular mechanisms underlying mitotic aneuploidy generation. This is in spite of findings that mitotic errors are equally or more prevalent than meiotic errors and arise independently of maternal age or fertility status (Vanneste et al. 2009; Chavez et al. 2012; McCoy et al. 2015a; McCoy et al. 2015b). Since the first three mitotic divisions are the most error-prone and activation of the embryonic genome does not occur until the 4- to 8-cell stage in the majority of mammals (Braude et al. 1988; Plante et al. 1994), it was suggested that maternally-inherited signaling factors regulating mitotic chromosome segregation may be lacking or compromised in early mammalian preimplantation embryos (Mantikou et al. 2012; Tsuiko et al. 2019).

There are several known contributors to aneuploidy and tumorigenesis in somatic cells, such as loss or prolonged chromosome cohesion, defective spindle attachments, abnormal centrosome number, and relaxed cell cycle checkpoints (Soto et al. 2019). Regardless of the mechanism, chromosomes that are mis-segregated during meiosis or mitosis will become encapsulated into micronuclei and can contribute to aneuploidy in subsequent divisions. In embryos, research has focused on the spindle assembly checkpoint (SAC) and primarily with mice that normally exhibit a low incidence of micronucleation and aneuploidy (Bolton et al. 2016; Treff et al. 2016; Vazquez-Diez et al. 2016). Thus, murine embryos are often treated with chemicals that inhibit spindle formation or SAC function to induce chromosome mis-segregation (Wei et al. 2011; Bolton et al. 2016; Vazquez-Diez et al. 2019; Singla et al. 2020), which target multiple genes and can have variable or off-target effects (Gascoigne and Taylor 2008; Miyazawa 2011). By monitoring bipolar attachment of spindle microtubules to kinetochores during mitosis, the mitotic checkpoint complex (MCC) prevents activation of the anaphase promoting complex/cyclosome (APC/C) and delays mitotic progression in the absence of stable bipolar kinetochore-microtubule attachments. This delay, however, is only temporary and cells with an unsatisfied checkpoint will eventually arrest or exit mitosis prematurely. The core components of the MCC are evolutionarily conserved and include CDC20, as well as the serine/threonine kinases, BUB1B, BUB3, and MAD2. BUB1B (also known as BUBR1), the largest of the MCC proteins, is normally present throughout the cell cycle and proposed to have both SAC-dependent and independent functions (Elowe et al. 2010). Besides being directly associated with unattached or incorrectly attached kinetochores, BUB1B also has a role in stabilizing kinetochore–microtubule attachments and chromosome alignment via BUB3 binding (Meraldi and Sorger 2005). Without BUB1B, the MCC no longer localizes to unattached kinetochores to prevent incorrect or deficient spindle attachments, resulting in the generation of aneuploid daughter cells (Lampson and Kapoor 2005). Whether the MCC is functional in the initial mitotic divisions of mammalian preimplantation development is currently unclear (Wei et al. 2011; Vazquez-Diez et al. 2019) and remains to be studied in a mammal that normally undergoes a high incidence of mitotic aneuploidy without the need for chemical induction.

Cattle are mono-ovulatory and share other key characteristics of preimplantation development with humans, including the timing of the first mitotic divisions, stage at which the major wave of embryonic genome activation (EGA) occurs, and approximate percentage of embryos that typically reach the blastocyst stage (Alper et al. 2001; Wong et al. 2010; Sugimura et al. 2012). Furthermore, single-nucleotide polymorphism (SNP) genotyping and next generation sequencing (NGS) revealed that the frequency of aneuploidy in cattle is likely similar to humans (Destouni et al. 2016; Hornak et al. 2016; Tsuiko et al. 2017). Destouni *et al*. also demonstrated that bovine zygotes can segregate parental genomes into different blastomeres during the first cleavage division, but the mechanism by which this occurs has not yet been determined (Destouni et al. 2016). Thus, with the ethical and technical limitations of human embryo research, bovine embryos represent a suitable model for studying the dynamics of micronuclei formation and aneuploidy generation during preimplantation development. In this study, we used a combination of time-lapse and live-cell fluorescent imaging with single-cell DNA-seq (scDNA-seq) for copy number variation (CNV) analysis, to assess mitotic divisions in bovine embryos from the zygote to 12-cell stage and visualize chromosome segregation in real-time. We also evaluated the lack of MCC function on cytokinesis, micronucleation, mitotic aneuploidy, and developmental arrest to determine if defective checkpoint signaling contributes to aneuploidy during early embryogenesis in higher-order mammals.

## RESULTS

### Micro- and multi-nucleation is common in early cleavage-stage bovine embryos

While micronuclei-like structures have been detected in bovine embryos previously (Yao et al. 2018), their prevalence or whether they were associated with a particular stage of preimplantation development was not determined. To address this, we generated a large number (N=53) of bovine embryos by *in vitro* fertilization (IVF) and fixed them at the zygote to blastocyst stage to evaluate DNA integrity with DAPI and nuclear structure by immunostaining for the nuclear envelope marker, LAMIN-B1 (LMNB1; **Fig. 1A**). Immunofluorescent labeling revealed the presence of micronuclei as early as the zygote stage that were distinct from the maternal and paternal pronuclei (**Fig. 1B**). Several micronuclei, as well as multiple nuclei (multi-nuclei) of similar size, were also detected at the 2- to 4-cell stage (**Fig. 1C**). Overall, ~37.7% (N=20/53) of early cleavage-stage embryos exhibited micro-/multi-nuclei formation in one or more blastomeres. This suggests that unlike mice, which rarely exhibit micronucleation during initial mitotic divisions (Vazquez-Diez et al. 2019), encapsulation of mis-segregated chromosomes into micronuclei prior to EGA is conserved between cattle and primates (Chavez et al. 2012; Daughtry et al. 2019). A similar examination of blastocysts also immunostained for the trophoblast marker, Caudal Type Homeobox 2 (CDX2), demonstrated that micronuclei often reside in the trophectoderm (TE; **Fig. 1D**), but can also be contained within the inner cell mass (ICM) of the embryo (**Fig. 1E**).

**Figure 1.**
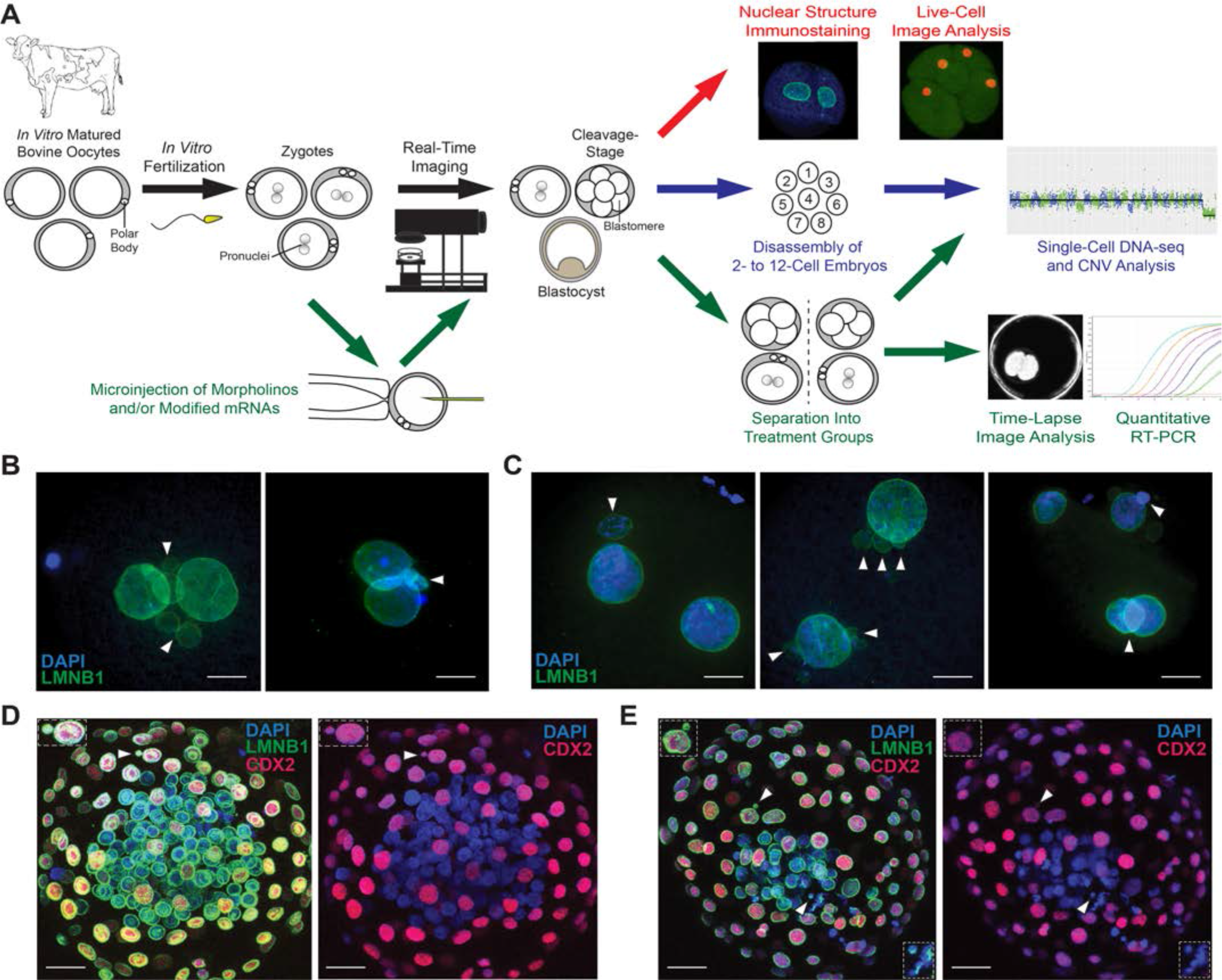
Investigating the dynamics of mitotic chromosome segregation and MCC fidelity in bovine embryos. **(A)** *In vitro* produced bovine oocytes underwent IVF and the resulting zygotes non-invasively monitored by time-lapse image analysis until collection for immunostaining of nuclear structure. Another subset of zygotes was microinjected with fluorescently labeled modified mRNAs and chromosome segregation visualized during the first three mitotic divisions in real-time by live-cell confocal microscopy. Cleavage-stage embryos were disassembled into single blastomeres at the 2- to 12-cell stage for scDNA-seq and CNV analysis to determine the precise frequency of aneuploidy at multiple cleavage stages. Other zygotes were microinjected with non-overlapping morpholinos targeting the mitotic checkpoint protein, BUB1B, and/or modified BUB1B mRNA to test the effect and specificity of MCC inhibition on chromosome segregation, division dynamics, and preimplantation development. Gene expression profiling was also conducted on a subset of MCC deficit zygotes versus controls by quantitative RT-PCR to identify changes in gene abundance and molecular pathways associated with BUB1B knockdown. **(B)** Immunostaining of zygotes and **(C)** cleavage-stage embryos with LMNB1 (green) using DAPI (blue) to visualize DNA revealed several micro- and multi-nuclei (white arrows). **(D)** Blastocysts also immunostained for the trophoblast marker, CDX2 (red), showed that micronuclei are often present in the TE, **(E)** but can also be retained within the ICM of the embryo. Scale bar = 10μm.

### Live-cell fluorescent imaging reveals micronuclei fate and origin of uniparental cells

To confirm the frequency of micro- and multi-nuclei in cleavage-stage embryos and determine the fate of these nuclear structures in real-time, we microinjected bovine zygotes (N=90) with fluorescently labeled modified mRNAs and monitored the first three mitotic divisions by live-cell confocal microscopy (**Fig. 1A**). While Histone H2B and/or LMNB1 were used to visualize DNA and nuclear envelope, respectively, F-actin was injected to distinguish blastomeres (**Supplemental Movie S1**). Of the microinjected embryos, ~18.9% (N=17/90) failed to complete cytokinesis during microscopic evaluation, whereas ~53.3% (N=49/90) exhibited normal bipolar divisions and ~27.8% (N=25/90) underwent multipolar divisions from 1- to 3-cells or more (**Fig. 2A**). In accordance with our immunostaining findings, ~31.1% (N=28/90) of the embryos contained micro- and/or multi-nuclei and anaphase lagging of chromosomes was detected prior to their formation in three of these embryos at the zygote (**Fig. 2B**) or 2-cell stage (**Fig. 2C**). Micro- and multi-nucleation was more frequently associated with bipolar divisions (**Fig. 2A**) and an examination of micronuclei fate demonstrated an equal incidence of unilateral inheritance (**Fig. 2D**) or fusion back with the primary nucleus (**Fig. 2E**), while a smaller percentage appeared to form a chromatin bridge with a neighboring blastomere (**Fig. 2F**, **Supplemental Fig. S1** and **Supplemental Movie S1**). Interestingly, the majority of multipolar embryos (76%; N=19/25) underwent an abnormal division after bypassing syngamy, or the fusion of maternal and paternal pronuclei (**Fig. 2G**), and/or produced daughter cells that did not contain any apparent nuclear structure (**Fig. 2H**). These results helped explain previous findings of blastomeres with uniparental origins and those that completely lacked nuclear DNA when assessed for CNV, respectively (Destouni et al. 2016; Ottolini et al. 2017; Daughtry et al. 2019; Middelkamp et al. 2020).

**Figure 2.**
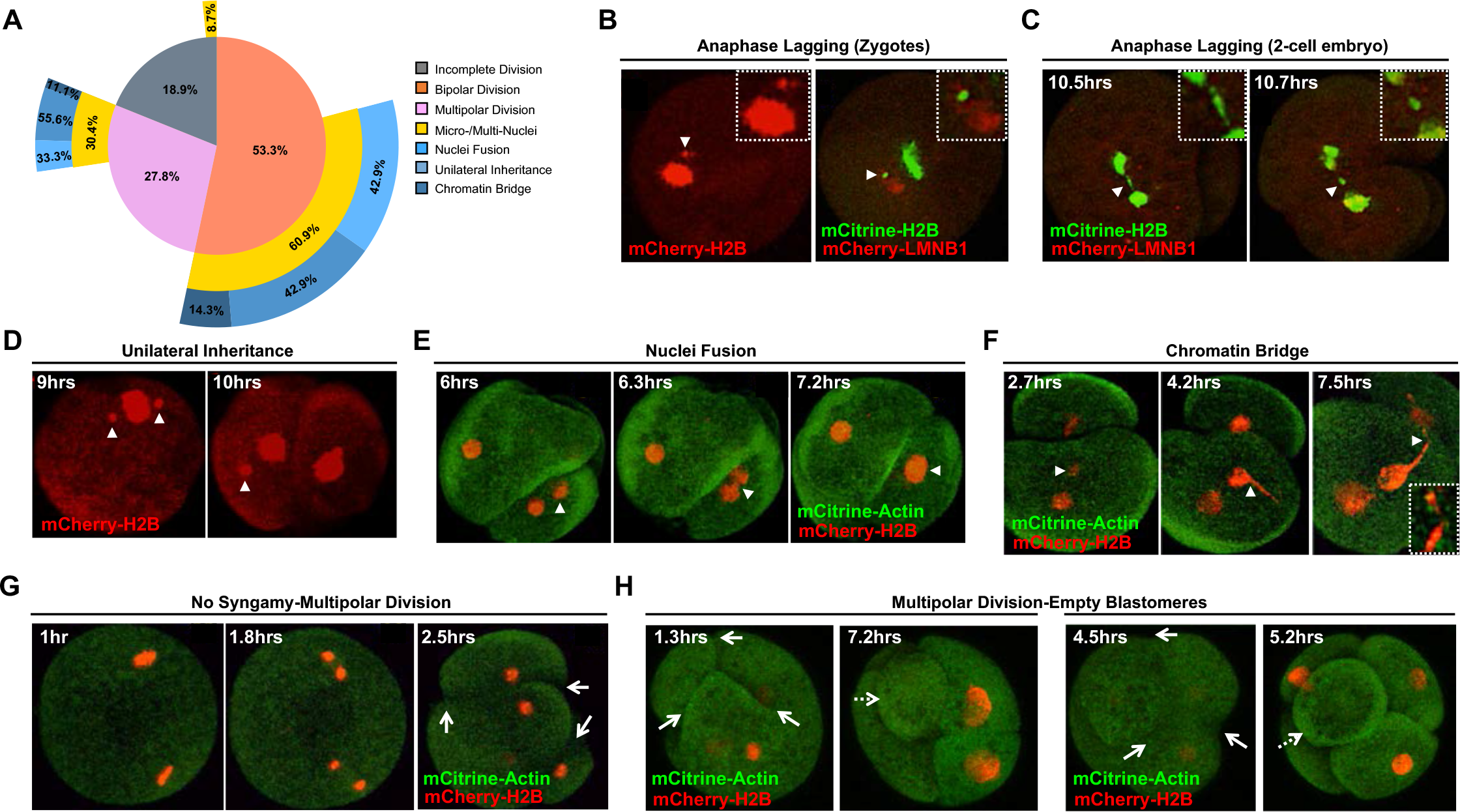
Live-cell fluorescent imaging reveals micronuclei fate and uniparental genome distribution to daughter cells. Bovine zygotes were microinjected with fluorescently labeled modified mRNAs (mCitrine or mCherry) to visualize DNA (Histone H2B) or nuclear structure (LMNB1) and distinguish blastomeres (FActin) by live-cell confocal microscopy during the first three mitotic divisions (N=90). **(A)** A Venn-Pie that shows the percentage of embryos that did not complete cytokinesis (gray), exhibited normal bipolar divisions (orange), or underwent multipolar divisions at the zygote or 2-cell stage (pink). The percentage of embryos with micro- and/or multi-nuclei (MN; yellow) associated with each type of division is also shown. Micronuclei fate is represented as those that formed a chromatin bridge (dark blue), exhibited unilateral inheritance (medium blue), or re-fused with the primary nucleus (light blue). Note that most embryos underwent bipolar divisions and were more likely to contain micronuclei than multipolar embryos. **(B)** Anaphase lagging of chromosomes (white arrowheads) was detected in certain embryos at the zygote or **(C)** 2-cell stage prior to micronuclei formation. **(D)** An examination of micronuclei fate demonstrated that a relatively equal proportion persist and undergo unilateral inheritance or **(E)** fuse back with the primary nucleus, **(F)** with a small number exhibiting what appeared to be a chromatin bridge between blastomeres following micronuclei formation (white arrowheads). **(G)** The majority of multipolar embryos (white solid arrows) bypassed pronuclear fusion (syngamy) prior to the abnormal division and **(H)** often produced blastomeres with uniparental origins and/or no apparent nuclear structure (white dashed arrows). Numbers in upper left corner represent the time in hours (hrs) since the start of imaging.

### Non-reciprocal mitotic errors and chaotic aneuploidy are prevalent in early cleavage divisions

Although SNP arrays or NGS have been used previously to assess aneuploidy in cleavage-stage bovine embryos, these studies reported a large range in aneuploidy frequency (~32-85%), examined a single stage of development, and/or evaluated only a portion of the embryo (Destouni et al. 2016; Hornak et al. 2016; Tsuiko et al. 2017). Therefore, our next objective was to determine the precise frequency of aneuploidy in a large number of bovine embryos (N=38) disassembled into individual cells at multiple cleavage stages using high-resolution scDNA-seq (**Fig. 1A** and **Supplemental Table S1**). All cells from the 38 embryos were assessed to ensure an accurate representation of the overall embryo, resulting in a total of 133 blastomeres analyzed from the 2- to 12-cell stage (**Fig. 3A**). Based on previously described criteria (Daughtry et al. 2019), we classified 25.6% (N=34/133) of blastomeres as euploid, 35.3% (N=47/133) as aneuploid, 3% (N=4/133) solely containing segmental errors, and 17.3% (N=23/133) exhibiting chaotic aneuploidy, with the remaining cells either failing WGA (10.5%; N=14/133) or identified as empty due to the amplification and detection of only mitochondrial DNA (8.3%; N=11/133). After reconstructing each embryo, we determined that ~16% (N=6/38) were entirely euploid, whereas ~55% (N=21/38) were comprised of only aneuploid cells (**Fig. 3B**). An additional ~29% (N=11/38) were categorized as mosaic since they contained a combination of both euploid and aneuploid blastomeres. Of the embryos with mosaicism, ~18% (N=2/11) had incurred segmental errors only, or DNA breaks of 15 Mb in length or larger that did not affect the whole chromosome. The X chromosome was by far the most frequently impacted by whole chromosomal losses and gains, whereas chromosome 5 (human chromosomes 12 and 22), 7 (human chromosomes 5 and 19), 11 (human chromosomes 3 and 9), and 29 (human chromosome 11) were commonly subjected to DNA breakage (**Fig. 3C**). While meiotic mis-segregation was identified in ~16% (N=6/38) of the embryos (**Fig. 3D**), mitotic aneuploidy accounted for the majority (~66%; N=25/38) of errors, with the remaining ~18% (N=7/38) exhibiting the genotypic complexity characteristic of chaotic aneuploidy (**Fig. 3E**). In addition, most (~67%; N=4/6) of the embryos with meiotic errors also experienced mitotic mis-segregation of different chromosomes than those originally affected during meiosis (**Fig. 3F**) and reciprocal losses and gains, whereby chromosomes lost from one blastomere are found in a sister blastomere, accounted for only ~25% (N=7/29) of the mitotic errors (**Fig. 3D** and **3F**).

**Figure 3.**
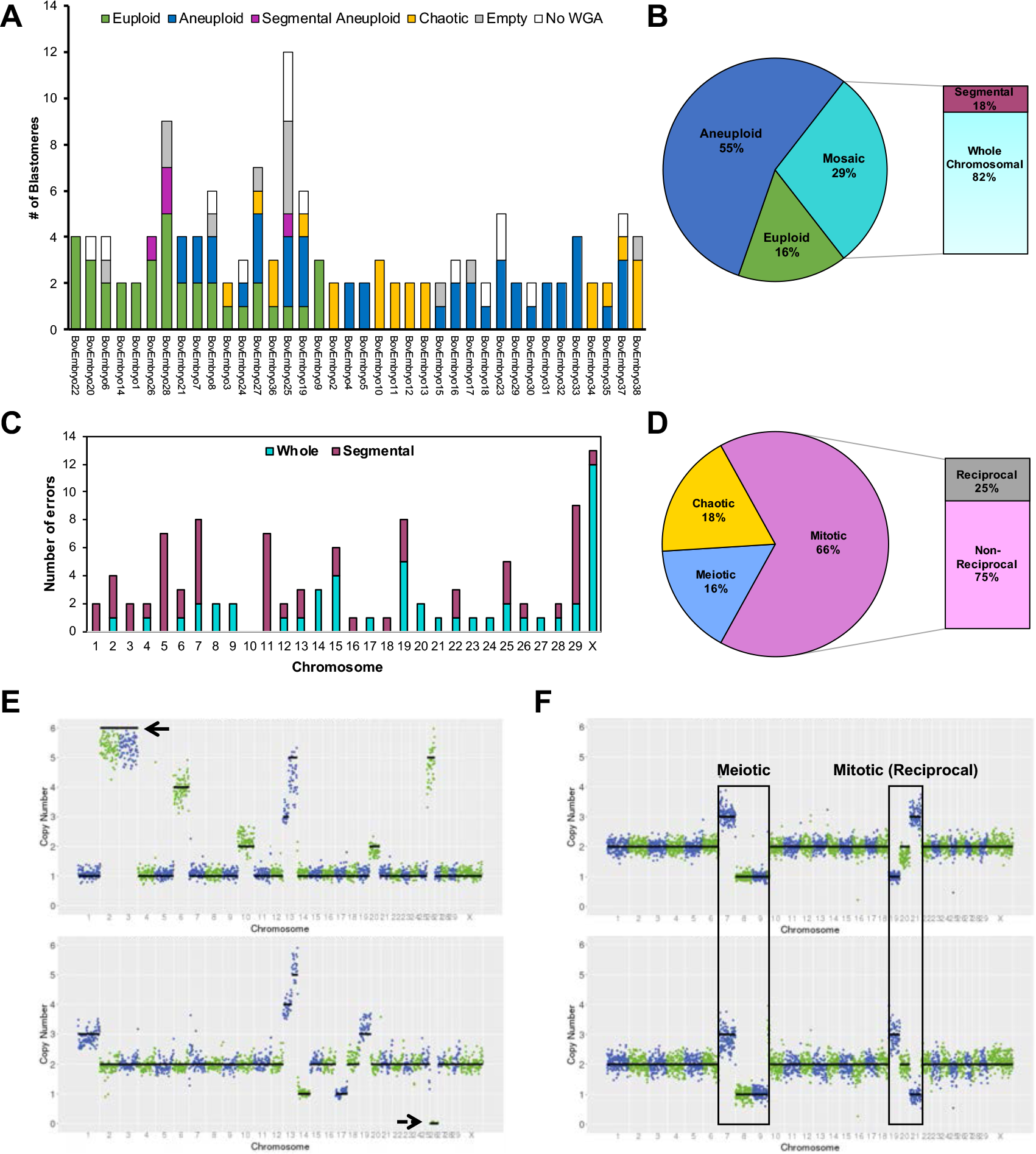
Comprehensive assessment of chromosomal abnormalities in early cleavage-stage embryos by scDNA-seq. **(A)** Whole chromosome and sub-chromosomal CNV was evaluated in bovine embryos from the 2- to 12-cell stage (N=38). Stacked bars represent all blastomeres (N=133) classified as euploid (green), aneuploid (blue), segmental aneuploid (purple), chaotic aneuploid (yellow), empty (grey) or failing to undergo WGA (white). **(B)** Pie chart showing the overall chromosome status of the embryos. **(C)** Number of whole or segmental chromosome losses and/or gains affecting each chromosome. Note the frequent mis-segregation of the X-chromosome and DNA breakage in chromosomes 5, 7, 11, and 29. **(D)** The percentage of aneuploid embryos with each type of chromosomal error. **(E)** CNV plots of blastomeres from two different embryos with chaotic aneuploidy showing up to 6 copies of certain chromosomes (top; black solid arrow) and a complete loss of other chromosomes (bottom; black dashed arrow). **(F)** Blastomeres from a 2-cell embryo with meiotic errors (Ch.7, 8, and 9) propagated during the first cleavage division that also experienced mitotic mis-segregation of different chromosomes (Ch.19 and 21) that were reciprocal.

### MCC deficiency induces atypical cytokinesis, blastomere asymmetry and embryo arrest

Since the chromosome constitution and division dynamics observed in certain embryos indicated deficient cell cycle checkpoints and there are conflicting reports on whether the MCC is functional at the early cleavage stage in mammals (Wei et al. 2011; Vazquez-Diez et al. 2019), our next objective was to determine if a lack of adequate checkpoints was associated with micronuclei formation and aneuploidy (**Fig. 1A**). Given negligible effects on mouse development from knockdown of another MCC component (Vazquez-Diez et al. 2019), we focused our attention on BUB1B/BUBR1, the largest of the MCC proteins that is present throughout the cell cycle (Elowe et al. 2010). Two non-overlapping morpholino antisense oligonucleotides (MAOs) were designed to specifically inhibit the translation of BUB1B mRNA by targeting the ATG translation start site (BUB1B MAO #1) or a sequence upstream within the 5’ UTR (BUB1B MAO #2) and tested before use in embryos (**Supplemental Fig. S2**). Zygotes were microinjected with either BUB1B MAO #1 (N=48), BUB1B MAO #2 (N=36), or standard control (Std Control) MAO (N=81) and cultured under a time-lapse imaging microscope to monitor developmental dynamics. Each embryo was morphologically assessed and categorized as having either normal or abnormal divisions for comparison to untreated (non-injected) embryos (N=180). In the BUB1B MAO #1 treatment group, 37.5% (N=18/48) of the zygotes failed to undergo the first cleavage division (**Table 1**) and a subset (8.3%; N=4/48) of these embryos attempted to divide by forming multiple cleavage furrows (**Fig. 4A**), but never successfully completed cytokinesis (**Supplemental Movie S2**). Of those BUB1B MAO #1 zygotes that divided, only a small proportion (18.8%; N=9/48) were normal bipolar divisions. Rather, many embryos (63.0%; N=17/27) exhibited abnormal cytokinesis, including multipolar divisions and/or blastomere asymmetry (**Supplemental Movie S3** and **Supplemental Movie S4**, respectively), with similar results obtained following injection with BUB1B MAO #2 (**Table 1** and **Fig. 4B**). Despite the phenotypic similarities between the two non-overlapping MAOs, we further assessed BUB1B MAO specificity by conducting embryo rescue experiments with modified BUB1B mRNA that would not be directly targeted by the MAO. BUB1B mRNA with a mutated MAO binding sequence was microinjected into zygotes, along with BUB1B MAO #1 (N=51), and embryos cultured up to the blastocyst stage (**Fig. 4C**). While no embryos formed blastocysts following injection of either the BUB1B MAO #1 or #2, 45% (N=23/51) of the BUB1B MAO #1+mRNA co-injected embryos underwent cleavage divisions and reached the blastocyst stage (**Fig. 4D**). This percentage was similar to that obtained from the non-injected embryos and following injection with the Std Control MAO, confirming that the knockdown of BUB1B expression and rescue of BUB1B-induced mitotic defects were specific.

**Table 1:** Division dynamics in untreated and MAO embryo treatment groups. Summary of the percentage of bovine zygotes that exhibited no division or attempted to divide as well as those that had normal bipolar/symmetric versus abnormal multipolar/asymmetric divisions following no treatment or microinjection with Std Control, BUB1B MAO #1, or BUB1B MAO #2. Attempted division was defined by the identification of cleavage furrows without the completion of cytokinesis. Note that in contrast to the controls, BUB1B MAO #1 and BUB1B MAO #2-injected embryos were more likely to undergo multipolar and/or asymmetric divisions.

|  | Untreated<br>(Non-Injected) | Std Control<br>MAO | BUB1B MAO<br>#1 | BUB1B<br>MAO #2 |
| --- | --- | --- | --- | --- |
| No<br>Division | 8.3%<br>(N=15/180) | 24.7%<br>(N=20/81) | 37.5%<br>(N=18/48) | 33.3%<br>(N=12/36) |
| Attempted<br>Division | 10%<br>(N=18/180) | 2.5%<br>(N=2/81) | 8.3%<br>(N=4/48) | 0.0%<br>(N=0/36) |
| Normal Bipolar/<br>Symmetric Division | 72.8%<br>(N=107/147) | 62.7%<br>(N=37/59) | 34.6%<br>(N=9/26) | 25.0%<br>(N=6/24) |
| Abnormal<br>Multipolar/<br>Asymmetric Division | 27.2%<br>(N=40/147) | 37.3%<br>(N=22/59) | 65.4%<br>(N=17/26) | 75.0%<br>(N=18/24) |
| Total Number of<br>Embryos | 180 | 81 | 48 | 36 |

**Figure 4.**
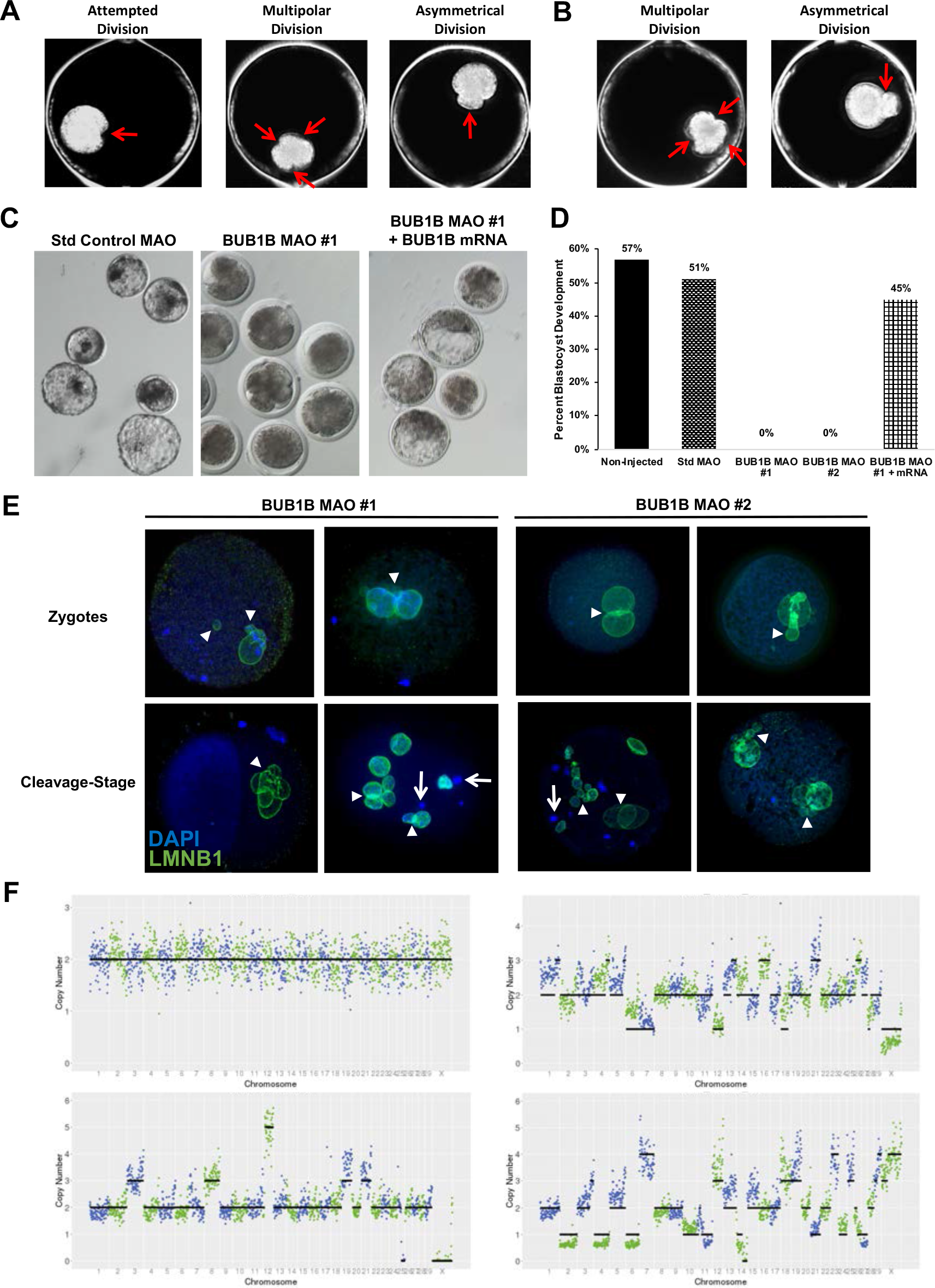
BUB1B knockdown induces multipolar divisions, chaotic aneuploidy, and developmental arrest. **(A)** Darkfield time-lapse imaging frames depicting the various embryo phenotypes (red arrows), including attempted division, multipolar division, and blastomere asymmetry observed following BUB1B MAO #1 or **(B)** BUB1B MAO #2 microinjection in bovine zygotes. **(C)** Representative stereomicroscope images of embryos and blastocysts from the Std control MAO, BUB1B MAO #1, and BUB1B MAO #1 plus BUB1B modified mRNA treatment groups. **(D)** Bar graph of the percentage of embryos that reached the blastocyst stage in non-injected, Std control MAO, BUB1B MAO #1, BUB1B MAO #2, or BUB1B MAO #1 plus BUB1B modified mRNA injected zygotes. While no blastocysts were obtained following BUB1B MAO #1 or #2 treatment, the co-injection of BUB1 MAO #1 and BUB1B modified mRNA was able to almost fully rescue the phenotype and restore blastocyst formation rates to that observed in controls. **(E)** Confocal images of LMNB1 (green) immunostaining in BUB1B MAO #1 or #2 treated embryos stained with DAPI (blue). Severely abnormal nuclear morphology and the presence of both micro- and multi-nuclei were detected (denoted with white arrowheads) in embryos at the zygote stage (top row) and cleavage-stage that exhibited abnormal cell divisions (bottom row). Note the DNA without nuclear envelope (white arrows) and the blastomere that completely lacked nuclear material in the 2-cell embryo located in the lower left image; Scale bars = 10μm. **(E)** CNV plots of blastomeres from different cleavage-stage embryos disassembled into single cells following BUB1B #1 MAO injection. While some euploid blastomeres were detected in BUB1B-injected embryos (upper left plot), most exhibited chaotic aneuploidy with multiple whole and sub-chromosomal losses and gains.

### MCC deficient embryos exhibit chaotic aneuploidy and asymmetric genome distribution

Because BUB1B MAO-injected embryos underwent atypical cytokinesis, we examined nuclear structure and CNV in MCC-deficient embryos by immunofluorescence and scDNA-seq, respectively (**Fig. 1A**). LMNB1 immunostaining revealed both micro- and multi-nuclei in BUB1B MAO #1 and #2 treated embryos that did not attempt division or were unable to complete the first cytokinesis (**Fig. 4E**). Similar abnormal nuclear structures, as well as empty blastomeres, were also observed in BUB1B MAO-injected embryos that successfully divided. Moreover, DNA that lacked or had defective nuclear envelope was also apparent in these MCC deficient embryos. Disassembly of the embryos into individual cells for assessment of DNA content and CNV analysis demonstrated that while some euploid blastomeres were obtained following BUB1B MAO injection, MCC deficiency mostly produced blastomeres with chaotic aneuploidy (**Fig. 4F**). Analogous to some of the non-injected controls (**Fig. 3E**), a complete loss of certain chromosomes and a gain of up to 5-6 copies of other chromosomes were detected, suggesting that the lack of MCC function permits premature mitotic exit and asymmetrical genome distribution in embryos.

### Lack of an intact MCC at the first division impacts cell cycle progression and kinase activity

Given that inappropriate expression of maternally-inherited signaling factors has been suggested to regulate early mitotic chromosome segregation in mammalian embryos (Mantikou et al. 2012; Tsuiko et al. 2019), we next determined whether MCC deficiency impacted the expression of other key developmental genes (**Fig. 1A**). Therefore, the relative abundance of maternal-effect, mitotic, cell cycle, EGA, and cell survival genes was assessed in individual BUB1B MAO #1 versus non-injected and Std Control-injected MAO embryos (**Supplemental Fig. S3** and **Supplemental Table S2**) via microfluidic quantitative RT-PCR (qRT-PCR). Besides *BUB1B*, other genes involved in cytokinesis and chromosome segregation such as amyloid beta precursor protein binding family B member 1 (*APBB1*), which inhibits cell cycle progression, aurora kinase B (*AURKB*), Polo-like kinase 1 (*PLK1*), and Ribosomal protein S6 kinase alpha-5 (*RPS6KA5*) were significantly downregulated in BUB1B MAO-injected embryos relative to the controls (**Fig. 5A**; p≤0.05). Additional genes, including those associated with the extracellular matrix (cartilage acidic protein 1; *CRTAC1* and ADAM metallopeptidase with thrombospondin type 1 motif 2; *ADAMTS2*) and stress response (Endoplasmic Reticulum Lectin 1; *ERLEC1*) were also significantly decreased in MCC deficient embryos in comparison to the non-injected and Std Control MAO-injected embryos. In contrast, genes involved in cell cycle progression such as Epithelial Cell Transforming 2 (*ECT2*), pogo transposable element derived with ZNF domain (*POGZ*), centromere protein F (*CENPF*), and Ribosomal protein S6 kinase alpha-4 (*RPS6KA4*), were significantly upregulated in BUB1B MAO-injected embryos, along with microtubule polymerization (HAUS augmin like complex subunit 6; *HAUS6*) or orientation (Synaptonemal complex protein 3; *SCP3*) genes (**Fig. 5B**; p≤0.05). Thus, in the absence of a functional MCC, we postulate that zygotes still entered mitosis, but were unable to obtain proper microtubule-kinetochore attachments prior to the first cytokinesis despite several attempts. This resulted in dysregulation of other kinases or cytoskeletal genes important for mitotic exit, cytokinesis, and chromosome segregation, confirming that MCC deficiency contributes to the large genotypic complexity observed during early cleavage divisions in higher-order mammals.

**Figure 5.**
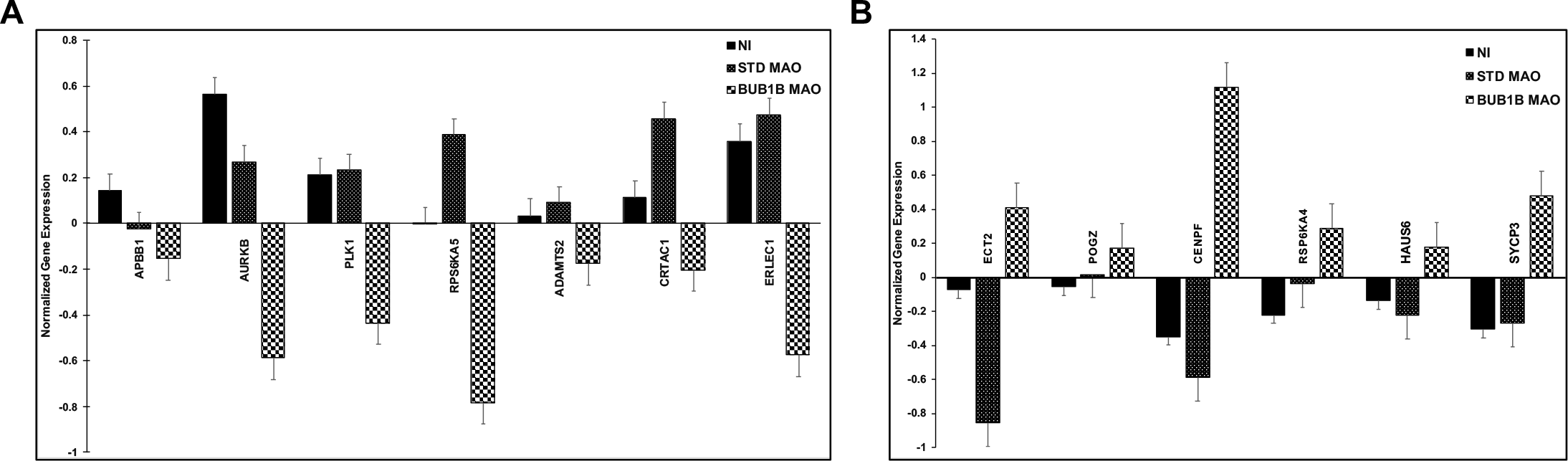
BUB1B deficiency in zygotes impacts the abundance of other cell cycle and mitosis-related genes. The relative abundance of several mitotic, cell cycle, developmentally-regulated, and cell survival genes was assessed via microfluidic quantitative RT-PCR (qRT-PCR) in non-injected (NI; N=5), Std Control MAO (N=5), and BUB1B MAO #1 (N=5) individual zygotes using gene-specific primers. **(A)** The genes that were significantly downregulated (p<0.05) in BUB1B MAO-injected embryos compared to the NI and Std Control MAO groups + standard error is shown in the bar graph. **(B)** A bar graph of the genes that were significantly upregulated in BUB1B MAO-injected embryos relative to the controls + standard error. CNRQ values of each gene was compared across embryo groups using the Mann-Whitney U-test. The full list of the 96 genes with primer sequences assessed by qRT-PCR is available in **Supplemental Fig. S3** and **Supplemental Table S2**, respectively.

## DISCUSSION

Aneuploidy is a major cause of embryo arrest, implantation failure, and spontaneous miscarriage across most mammalian species and yet, relatively little is still known about the molecular mechanism(s) underlying aneuploidy generation and pregnancy loss. While some understanding of mitotic mis-segregation derives from dividing somatic cells, the first embryonic cleavage divisions are fundamentally different since almost all of the mRNAs and proteins required for cytokinesis and chromosome segregation are maternally-inherited (Mantikou et al. 2012; Tsuiko et al. 2019). In addition, unlike tumors and cancer cells, which often overexpress cell cycle checkpoints and rarely sustain SAC gene mutations (Schvartzman et al. 2010), cleavage-stage human embryos have been shown to underexpress checkpoints and overexpress cell cycle drivers (Kiessling et al. 2010). Knockdown of a specific MCC component in mouse zygotes, however, had no effect on early cleavage divisions when mitotic aneuploidy typically occurs in other mammals (Vazquez-Diez et al. 2019). Moreover, because these studies were conducted with mice, which naturally exhibit a low incidence (~1-4%) of aneuploidy, the embryos were treated with chemicals to induce chromosome mis-segregation (Wei et al. 2011; Bolton et al. 2016; Treff et al. 2016; Vazquez-Diez et al. 2019; Singla et al. 2020). Using a combination of live-cell imaging, scDNA-seq, and genetic manipulation, we visualized mitotic chromosome segregation in real-time from the zygote to the ~12-cell stage and assessed the role of the MCC in embryos from an animal model that normally suffers from a comparable incidence of aneuploidy and developmental arrest as humans.

To determine the prevalence of chromosome mis-segregation in initial mitotic divisions, we first assessed the frequency of micronucleation throughout bovine preimplantation development. Of the cleavage-stage embryos examined by immunostaining or live-cell imaging, over ~30% contained micro- or multi-nuclei and anaphase lagging of chromosomes was detected in certain embryos prior to micronuclei formation. When we evaluated the cellular behaviors that might indicate how these atypical nuclear structures formed, we determined that most micronuclei-containing embryos underwent normal bipolar divisions, excluding abnormal cytokinesis as the primary mechanism. However, multipolar divisions were associated with a lack of syngamy and often produced cells that did not contain any apparent nuclear structure (**Fig. 6A**). In contrast to mouse embryos, which sustain spatial separation of parental genomes by dual-spindle formation (Mayer et al. 2000; Reichmann et al. 2018), embryos from other mammals are still thought to exhibit syngamy at the zygote stage (Kai et al. 2018; Yao et al. 2018). By avoiding syngamy and undergoing multipolar cytokinesis, zygotes differentially segregate entire parental genomes to daughter cells, providing a mechanism for previous findings of blastomeres with uniparental origins in both cattle and primates (Destouni et al. 2016; Ottolini et al. 2017; Daughtry et al. 2019; Middelkamp et al. 2020).

**Figure 6.**
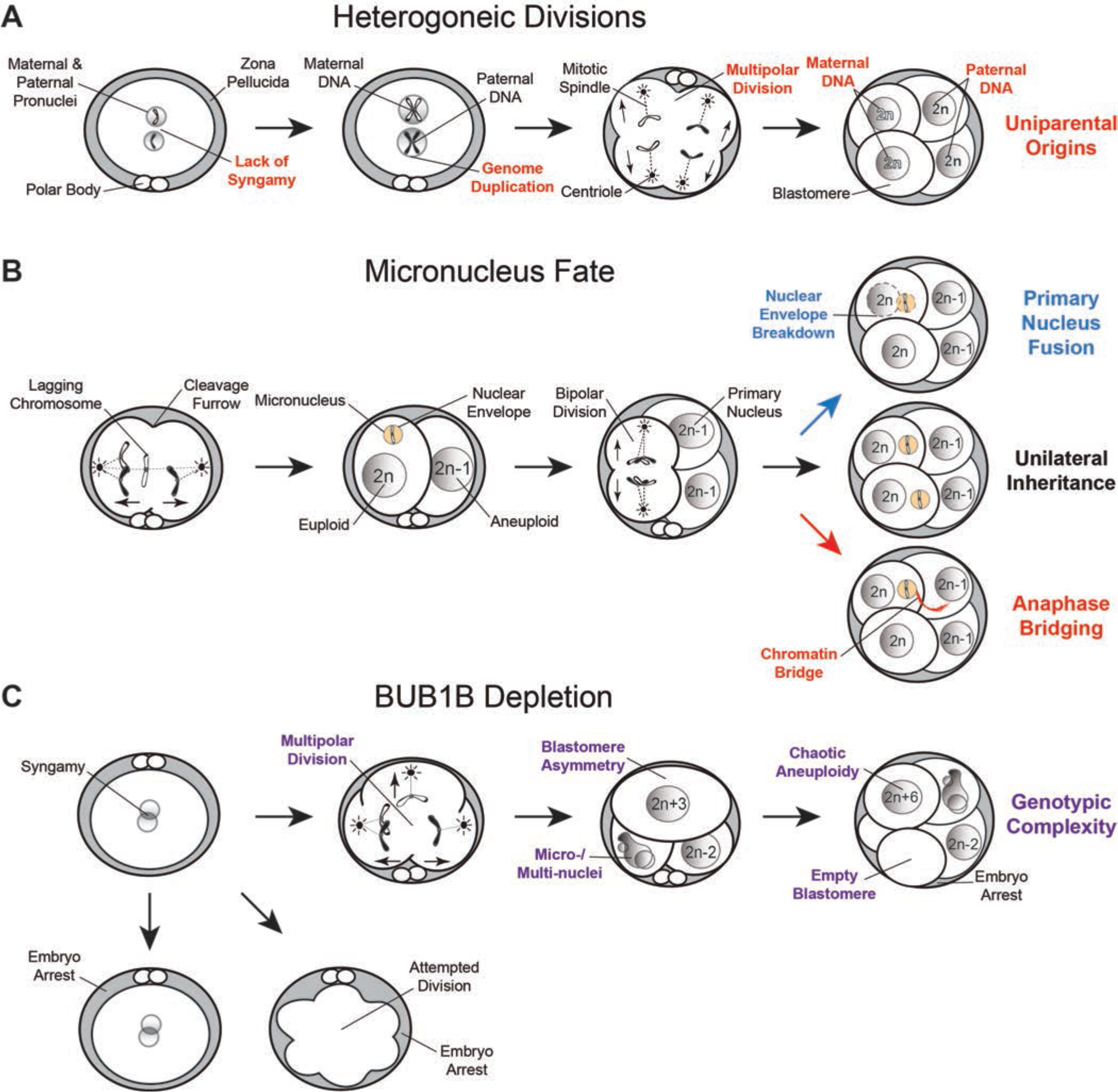
Summary of the major findings from the imaging, scDNA-seq, and gene knockdown studies. **(A)** Simplified model of how the lack of maternal and paternal pronuclear fusion (syngamy) at the zygote stage, followed by genome duplication and multipolar divisions, contributes to blastomeres with uniparental origins, or those that only contain maternal or paternal DNA. **(B)** Live-cell imaging also revealed the formation of anaphase lagging chromosomes likely from merotelic attachments prior to or during the first mitotic division. The chromosome(s) become encapsulated in nuclear envelope to form a micronucleus and the embryo continues to divide normally. In these subsequent bipolar divisions, most micronuclei either fuse back with the primary nucleus upon nuclear envelope breakdown or persist and undergo unilateral inheritance, but some micronuclei form a chromatin bridge with the nucleus of another blastomere during anaphase. **(C)** The depletion of BUB1B in zygotes resulted in no division or attempted division and embryo arrest, while multipolar divisions, blastomere asymmetry, and micro-/multi-nuclei were observed in MCC-deficient embryos that completed the first cytokinesis. These abnormal divisions also produced daughter cells with chaotic aneuploidy and/or empty blastomeres with no nuclear structure that induced embryo arrest and suggested that the lack of MCC permits the genotypic complexity detected at the early cleavage-stages of preimplantation development.

Examination of micronuclei fate in subsequent divisions revealed an equal incidence of unilateral inheritance and fusion back with the primary nucleus, with a smaller percentage of embryos exhibiting a chromatin bridge between blastomeres following micronuclei formation (**Fig. 6B**). Because cancer cell micronuclei have been shown to endure extensive DNA damage upon refusion with the primary nucleus (Crasta et al. 2012; Zhang et al. 2015), chromosomal integrity and the effects on developmental outcome will likely depend on which of these events occur. The significance of the chromatin bridging and whether it exacerbates aneuploidy or restores euploidy is unknown, but we suspect that the exchange of genetic material contributes to the large genotypic complexity reported in IVF embryos (Vanneste et al. 2009; Chavez et al. 2012; McCoy et al. 2015a; Daughtry et al. 2019). A similar assessment of bovine blastocysts determined that micronuclei often reside in the placental-derived TE, but can also be contained within the ICM of the embryo. While the presence of micronuclei in the ICM may be more detrimental, a recent study reported that there is no significant enrichment of aneuploid cells between the TE and ICM in human blastocysts (Starostik et al. 2020). Thus, micronuclei formation at this stage of development is probably more tolerated due to increased cell number or its impact on overall ploidy is not apparent until the postimplantation stages, which warrants further investigation.

Given the large aneuploidy range reported in previous bovine studies (Destouni et al. 2016; Hornak et al. 2016; Tsuiko et al. 2017), as well as differences in the stage or proportion of the embryo analyzed and methods used, we sought to comprehensively assess aneuploidy in all cells of embryos at multiple cleavage stages. After reconstructing each embryo and combining the results, we determined that ~55% of the embryos contained only aneuploid cells, whereas another ~29% were mosaic, all of which were primarily the product of non-reciprocal mitotic errors. In those embryos with meiotic errors, most also experienced mitotic mis-segregation of different chromosomes than those originally affected during meiosis. The remaining aneuploid embryos exhibited a compete loss and/or a gain of up to 6 copies of chromosomes characteristic of chaotic aneuploidy. This indicates that embryos with meiotic mis-segregation are more prone to mitotic errors and further propagated by subsequent divisions, further explaining the genotypic complexity observed in IVF embryos (Vanneste et al. 2009; Chavez et al. 2012; McCoy et al. 2015a; Daughtry et al. 2019).

Because of the apparent disparity on whether the MCC is functional in the early cleavage divisions of mammalian embryogenesis in previous studies (Wei et al. 2011; Vazquez-Diez et al. 2019), we investigated the consequences of MCC inhibition by directly targeting BUB1B in bovine zygotes. Following injection, BUB1B MAO embryos either failed to divide even after several attempts or exhibited abnormal divisions that were multipolar and/or asymmetrical (**Fig. 6C**). Furthermore, immunostaining of the BUB1B MAO treated embryos that did divide revealed blastomeres with severely abnormal nuclear structures or those that were completely devoid of DNA. CNV analysis of blastomeres that contained nuclear DNA showed a predominance of chaotic aneuploidy, with a complete loss or excessive number of chromosomal copies as described in some non-injected embryos and reported in primate embryos with multipolar divisions (Ottolini et al. 2017; Daughtry et al. 2019). Without BUB1B, we speculate that embryos were unable to obtain proper microtubule-kinetochore attachments prior to the first cytokinesis, resulting in failed MCC and arrest, or premature cell division and chromosome mis-segregation due to MCC dysregulation. The role of another MCC protein, Mad2, was also recently investigated in mouse embryos and while 40% Mad2 knockdown had no effect on blastocyst formation, it did double the number of micronuclei present at the morula stage (Vazquez-Diez et al. 2019). Both MAD2 and BUB1B bind CDC20 to prevent activation of the APC, but *in vitro* binding assays demonstrated that BUB1B is 12 times more effective than MAD2 in inhibiting CDC20 (Fang 2002). In addition, it was shown in *Drosophila* that the recruitment of CDC20 to the kinetochore requires BUB1B and not MAD2 (Li et al. 2010) and that BUB1B is maternally inherited (Perez-Mongiovi et al. 2005). Thus, these studies help explain the robust effect of BUB1B deficiency observed here and suggests that inhibition of the MCC via BUB1B knockdown impacts early cleavage divisions in higher-order mammals by allowing multipolar cytokinesis and asymmetrical genome partitioning to occur.

The expression of additional genes involved in mitosis and cell cycle progression was also affected by MCC inhibition and indicates that their abundance may be regulated by BUB1B availability in embryos. One of the downregulated genes included *Plk1*, which is conserved across both mammalian and non-mammalian species and has been shown to be important for the first mitosis in mouse zygotes (Baran et al. 2016). In somatic cells, PLK1 localization to non-attached kinetochores is required for the phosphorylation of BUB1B (Elowe et al. 2007) and promotes the interaction of BUB1B with phosphatases that, in turn, inhibit excessive aurora kinase activity at kinetochores through positive feedback (Suijkerbuijk et al. 2012). Therefore, the removal of BUB1B or inhibition of PLK1 increases the phosphorylation of kinase substrates, which has been shown to include *ECT2, POGZ*, and *HAUS6* (Kettenbach et al. 2011; Bibi et al. 2013), genes identified as upregulated following BUB1B knockdown here. Since BUB1B MAO-injected embryos also exhibited increased expression of *CENP-F* and *SYCP3* and both are regulated by PLK1 phosphorylation in other contexts (Santamaria et al. 2011), we suspect that these genes also serve as kinase substrates important for mitotic progression during embryogenesis. Additionally, we note that common maternal genotype variants spanning *PLK4*, another polo-like kinase family member, has been reported to play a role in tripolar divisions and aneuploidy in human embryos (McCoy et al. 2015a; McCoy et al. 2018). Thus, BUB1B likely cooperates with this regulatory network of kinases and their substrates to reinforce MCC function and ensure chromosome fidelity in embryos. Collectively, our results confirm a role for the MCC in maintaining proper chromosome segregation in initial cleavage divisions and show that the genotypic complexity observed in preimplantation embryos from higher-order mammals is likely contributed by deficiency in BUB1B and/or other maternally-inherited factors.

## MATERIALS AND METHODS

### Experimental design

Using a combination of live-cell imaging, scDNA-seq for CNV analysis, and genetic manipulation of embryos, we developed an experimental approach to assess mitotic divisions and chromosome segregation throughout bovine preimplantation development (**Fig. 1A**). First, we fertilized mature oocytes, cultured resultant zygotes under a time-lapse imaging microscope to monitor embryo developmental dynamics, and evaluated DNA integrity and nuclear structure by immunofluorescence up to blastocyst stage (N=53). We confirmed our findings by live-cell confocal microscopy of zygotes microinjected with fluorescently-labeled modified mRNAs and visualization of the initial mitotic divisions in real-time (N=90). Cleavage-stage embryos between 2- and 12-cells were then disassembled into single blastomeres for comprehensive assessment of meiotic and/or mitotic errors (N=38). Lastly, the role of the MCC in aneuploidy generation was determined by microinjecting zygotes with BUB1B MAOs (N=84) or a Std Control MAO (N=81) for comparison to non-injected embryos (N=180) and embryos co-injected with BUB1B MAO and BUB1B modified mRNA (N=85) by time-lapse monitoring, immunostaining, CNV analysis, and/or microfluidic quantitative RT-PCR.

### Reagents and media

All chemicals were obtained from Sigma-Aldrich (St. Louis, MO, USA) or Fisher Scientific (Pittsburgh, PA, USA) unless otherwise stated. Tyrode’s albumin lactate pyruvate (TALP) medium with Hepes (TALP-Hepes) was used as washing media and contained 114mM NaCL, 3.2mM KCl, 25mM NaHCO_3_, 0.34mM NaH_2_PO_4_-H_2_O, 10mM C_3_H_5_NaO_3_, 2mM CaCl_2_-H_2_O, 0.5mM MgCl_2_-6H_2_O, 10.9 mM Hepes, 0.25mM sodium pyruvate, 1μl/ml Phenol Red, 3mg/ml FAF-BSA, 100μM Gentamicin Sulfate. For fertilization, TALP-IVF was used and comprised of 114mM NaCL, 3.2mM KCl, 25mM NAHCO_3_, 0.34mM NaH_2_PO_4_-H_2_O, 10mM C_3_H_5_NaO_3_, 2mM CaCl_2_-H_2_O, 0.5mM MgCl_2_-6H_2_O, 1μl/ml Phenol Red, 0.25mM sodium pyruvate, 100units/ml penicillin, 100μg/ml streptomycin, 1μM epinephrine, 0.02 mM penicillamine, 10μM hypotaurine, 6mg/ml FAF-BSA, and 10mg/ml heparin.

### IVF and embryo culture

Cumulus-oocyte complexes (COC) were retrieved by follicular aspiration of ovaries collected at a commercial abattoir (DeSoto Biosciences, Seymour, TN, USA). Those COCs with at least three layers of compact cumulus cells and homogeneous cytoplasm were placed in groups of 50 in 2ml sterile glass vials containing 1ml of oocyte maturation medium, covered with mineral oil, and equilibrated in 5% CO_2_. Tubes with COCs were shipped overnight in a portable incubator (Minitube USA Inc., Verona, WI, USA) at 38.5°C. Following 24h of maturation, COCs were washed 3 times in TALP-Hepes followed by a final wash in fertilization media, before placement in a 4-well dish (Nunc™; Fisher Scientific) containing 0.5ml of fertilization media. Semen from either Racer (014HO07296) from Accelerated Genetics (Baraboo, WI, USA) or Colt P-red (7HO10904) from Select Sires (Plain City, OH, USA was obtained for IVF. Sperm were purified from frozen-thawed straws using a gradient [50% (v/v) and 90% (v/v)] of Isolate (Irvine Scientific, Santa Ana, CA), washed two times in fertilization media by centrifugation at 100 RCF, and diluted to a final concentration of 1 million/ml in the fertilization dish. Fertilization was allowed to commence for 17–19 h at 38.5°C in a humidified atmosphere of 5% CO_2_. Zygotes were denuded from the surrounding cumulus cells by vortexing for 4 min in 200μl of TALP-Hepes with 0.5% (w/v) hyaluronidase (Sigma-Aldrich) and washed in fresh TALP-Hepes.

### Time-lapse imaging

Denuded zygotes were transferred to custom Eeva™ 12-well polystyrene dishes (Progyny, Inc., New York, NY; formerly Auxogyn, Inc.) containing 100μl drops of BO-IVC culture media (IVF Bioscience; Falmouth, Cornwall, UK) under mineral oil (CooperSurgical, Trumbull, CT) and cultured at 38.5°C in a humidified atmosphere of 5% CO_2_, 5% O_2_, and 90% N_2_. Embryos were monitored with an Eeva™ darkfield 2.2.1 or bimodal (darkfield/brightfield) 2.3.5 time-lapse microscope system (Progyny, Inc) housed in a small tri-gas incubator (Panasonic Healthcare, Japan) as previously described (Vera-Rodriguez et al. 2015). Images were taken every 5 min with a 0.6 second exposure time. Each image was time stamped with a frame number and all images compiled into an AVI movie using FIJI software (NIH, Bethesda, MD) version 2.0.0 (Schindelin et al. 2012) for assessment of mitotic divisions by two independent reviewers.

### Immunofluorescent labeling

Embryos were washed in PBS with 0.1% BSA and 0.1% Tween-20 (PBST; Calbiochem, San Diego, CA) and fixed with 4% paraformaldehyde (Alfa Aesar, Ward Hill, MA) in PBST for 20 min. at room temperature (RT). Once fixed, the embryos were washed with gentle shaking three times for a total of 15 min. in PBS-T to remove residual fixative. Embryos were permeabilized in 1% Triton-X (Calbiochem) for one hour at RT and washed in PBST as described above. To block non-specific antibody binding, embryos were transferred to a 7% donkey serum (Jackson ImmunoResearch Laboratories, Inc., West Grove, PA)/PBS-T solution for either 1 hour at RT or overnight at 4°C. An antibody against LMNB1 (catalog #ab16048, Abcam, Cambridge, MA) was diluted 1:1,000, while the CDX2 mouse monoclonal antibody (clone #CDX2-88, Abcam) was diluted 1:100 in PBS-T with 1% donkey serum, and embryos stained for 1 hour at RT or overnight at 4°C. Primary LMNB1 and CDX2 immunosignals were detected using 488-conjugated donkey anti-rabbit or 647-conjugated donkey anti-mouse Alexa Fluor secondary antibodies (Thermo Fisher), respectively, at a 1:250 dilution with 1% donkey serum in PBS-T at RT for 1 hour in the dark. Embryos were washed in PBS-T and the DNA stained with 1μg/ml DAPI for 15 min. Embryos were mounted on slides using Prolong Diamond mounting medium (Invitrogen, Carlsbad, CA, USA). Immunofluorescence was initially visualized on a Nikon Eclipse Ti-U fluorescent microscope system and images captured using a Nikon DS-Ri2 color camera and confirmed with a Leica SP5 AOBS spectral confocal system. Z-stacks, 1–5uM apart, were imaged one fluorophore at a time to avoid spectral overlap between channels. Stacked images and individual channels for each color were combined into composite images using FIJI software version 2.0.0.

### Modified mRNA construction

Plasmids containing the coding sequence (CDS) for mCitrine-Lifeact (Addgene #54733), which labels filamentous actin (F-actin), mCherry-Histone H2B-C-10 (Addgene #55057), and mCherry-LAMINB1-10 (Plasmid #55069) were a gift from Dr. Michael Davidson’s laboratory and deposited in Addgene (Cambridge, MA). Custom primers containing a 5’-T7 promoter sequence were used to amplify each fluorescent tag-mRNA fusion construct as follows:

T7_mCitrine_F: CTAGCTTAATACGACTCACTATAGGGCGGTCGCCACCATGGTGA
LifeAct_R: TTACTTGTACAGCTCGTCCATGCCGAGAGTGATCCCGGC
T7_mCherry_F: AATTAATACGACTCACTATAGGGAGAGCCACCATGGTGAGCAA
H2B_R: GCGGCCGCTTTACTTGT
LAMINB1_R: TCCGGTGGATCCCTACATAA

PCR amplification was performed with high fidelity Platinum Taq polymerase (Thermo Fisher) under the following conditions: 94°C for 2 min., followed by 35 cycles of 94°C-30 sec., 70°C-30 sec. and, 72°C-3 min. PCR products were purified with the QIAquick PCR Purification kit (Qiagen; Hilden, Germany), then underwent *in vitro* transcription using the mMessage Machine T7 Transcription Kit (Invitrogen). Following the synthesis of capped mRNA, the MEGAclear transcription clean up kit (Invitrogen) was used to purify and concentrate the final modified mRNA product.

### Live-cell imaging

Bovine zygotes were microinjected with mCitrine-Lifeact and either mCherry-H2B or mCherry-LAMINB1 mRNAs at a concentration of 20 ng/ul each in the presence of Alexa Fluor 488 labeled Dextran (Invitrogen) using a CellTram vario, electronic microinjector and Transferman NK 2 Micromanipulators (Eppendorf, Hauppauge, New York, USA). Zygotes that exhibited mCherry fluorescent signal within 4-6 hours following microinjection were selected for overnight imaging. Imaging dishes were prepared by placing 20ul drops of BO-IVC media on glass bottom dishes (Matek Corporation; Ashland, MA) and covering with mineral oil. A Zeiss LSM 880 laser-scanning confocal microscope with 10X objective and Fast Airy capabilities was used to capture fluorescent images of embryos for 18-20 hours, which encompassed the first three mitotic divisions. Z-stack images were taken every 1.5μm for a total of ~60 slices covering a 90um range at 10 min. intervals. Each fluorophore was acquired independently to prevent crosstalk and maximize scanning speed. Individual images underwent Airyscan processing using Zeiss software and were compiled into videos with individual embryo labels using FIJI. Assessment of cytoplasmic and nuclear structure in embryos during mitotic divisions was completed by two independent reviewers.

### Embryo disassembly

Embryos were disassembled under a stereomicroscope equipped with a heated stage and digital camera (Leica Microsystems, Buffalo Grove, IL) for documentation. The zona pellucida was removed from each embryo by a 30 second exposure to warm Acidified Tyrode’s Solution (EMD Millipore, Temecula, CA), followed by 30-60 seconds in 0.1% (w/v) pronase (Sigma, St. Louis, MO, USA). Once ZP free, embryos were washed in TALP-Hepes and gently manipulated using a STRIPPER pipettor (Origio, Målov, Denmark), with or without brief exposure to warm 0.05% trypsin-EDTA (Thermo Fisher Scientific, Waltham, MA) as necessary, until all blastomeres were separated. Following disassembly, each blastomere and cellular fragment if present was washed three times with Ca^2+^ and Mg^2+^-free PBS (Fisher Scientific), collected into individual PCR tubes in ~2μL of PBS, and snap frozen on dry ice. Downstream analysis was completed only for embryos where the disassembly process was successful for all blastomeres.

### DNA library preparation

Single blastomeres and cellular fragments underwent DNA extraction and WGA using the PicoPLEX single-cell WGA Kit (Rubicon Genomics, Ann Arbor, MI) according to the manufacturer’s instructions with slight modifications. Cells were lysed at 75°C for 10 min. followed by pre-amplification at 95°C for 2 min. and 12 cycles of gradient PCR with PicoPLEX pre-amp enzyme and primer mix. Pre-amplified DNA was further amplified with PicoPLEX amplification enzyme and 48 uniquely-indexed Illumina sequencing adapters provided by the kit or custom adapters with indices designed as previously described (Vitak et al. 2017; Daughtry et al. 2019). Adapter PCR amplification consisted of a 95°C hotstart for 4 min., four cycles of 95°C for 20 sec., 63°C for 25 sec., and 72°C for 40 sec. and seven cycles of 95°C for 20 sec. and 72°C for 55 sec. Libraries were quantified with a Qubit High Sensitivity (HS) DNA assay (Life Technologies, Carlsbad, CA). Amplified DNA from each blastomere (50ng) and cellular fragment (25ng) was pooled and purified with AMPure^®^ XP beads (Beckman Coulter, Indianapolis, IN). Final library quality assessment was performed on a 2200 TapeStation (Agilent, Santa Clara, CA).

### Multiplex scDNA-seq

Pooled libraries were sequenced on an Illumina NextSeq 500 using a 75-cycle kit with a modified single-end workflow that incorporated 14 dark cycles at the start of the first read prior to the imaged cycles. This step excluded the quasi-random priming sequences that are G-rich and lack a fluorophore for the two-color chemistry utilized by the NextSeq platform during cluster assignment. A total of ~3.5×10^6^ reads/sample were generated. All raw sample reads were demultiplexed and sequencing quality assessed with FastQC (Krueger et al. 2011). Illumina adapters were removed from raw reads with the sequence grooming tool, Cutadapt (Chen et al. 2014), which trimmed 15 bases on the 5’ end and five bases from the 3’ end, resulting in reads of 120 bp on average. Trimmed reads were aligned to the most recent bovine reference genome, BosTau8 (Zimin et al. 2009), using the BWA-MEM option of the Burrows-Wheeler Alignment Tool with default alignment parameters (Salavert Torres J and J 2012). Resulting bam files were filtered to remove alignments with quality scores below 30 (Q<30) as well as alignment duplicates that were likely the result of PCR artifacts with the Samtools suite (Ramirez-Gonzalez et al. 2012). The average number of filtered and uniquely mapped sequencing reads in individual libraries was between 1.9 and 2.2 million.

### CNV analysis

CNV was determined by the integration of two previously developed bioinformatics pipelines, Variable Non-Overlapping Window Circular Binary Segmentation (VNOWC) and the Circular Binary Segmentation/Hidden Markov Model (CBS/HMM) Intersect termed CHI, as previously described (Vitak et al. 2017; Daughtry et al. 2019). All CNV calls from the two pipelines generated profiles of variable sized windows that were intersected on a window-by-window basis. Because other low-input sequencing studies have shown that CNV can be reliably assessed at a 15 Mb resolution with 0.5-1X genome coverage (Lee et al. 2013; Zhou et al. 2018), we classified breaks of 15 Mb in length or larger that did not affect the whole chromosome as segmental. Only whole and segmental CNV calls in agreement between the VNOWC and CHI methods at window sizes containing 4,000 reads were considered. Chaotic aneuploidy was classified by the loss or gain of greater than four whole and/or broken chromosomes as previously described (Daughtry et al. 2019). Additional classification of each aneuploidy as meiotic or mitotic in origin was accomplished by determining whether a loss or gain of the same chromosome was detected in all blastomeres (meiotic) or if different and/or reciprocal chromosome losses and gains were observed between blastomeres (mitotic).

### MAO Design

Two non-overlapping MAOs were designed and synthesized by Gene Tools (Philomath, OR) to specifically target bovine BUB1B (Ensembl transcript ID: ENSBTAT00000009521.5). BUB1B MAO #1 (TTTCCTTCTGCATCGCCGCCATC) specifically targeted the ATG start codon of the BUB1B mRNA coding sequence, while BUB1B MAO #2 (CGATCTGAGGCTCTGAAGAAAGGCC) targeted upstream of MAO #1 in the 5’ UTR of bovine *BUB1B*. A Std Control MO (CCTCTTACCTCAGTTACAATTTATA) that targets a splice site mutant of the human hemoglobin beta-chain (HBB) gene (GenBank accession no. AY605051) that is not present in the *Bos Taurus* genome served as a control. Both BUB1B and Std Control MAO where synthesized with a 3’-carboxyfluorescein tag to aid in visualization during cell transfection and embryo manipulation.

### Assessment of BUB1B MAO efficiency

Before use in embryos, the BUB1B MAOs were first tested using the Madin-Darby Bovine Kidney (MDBK) epithelial cell line (Madin and Darby 1958). MDBK cells were plated on poly-L-lysine treated coverslips, and grown to 70% confluency prior to MAO treatment. The cells were incubated with 6μl/ml Endo-Porter delivery reagent containing DMSO (Gene Tools) and 2, 4, or 8μM of either BUB1B MAO #1 or Std control MAO and cultured in Eagle’s Minimum Essential Medium modified to contain Earle’s Balanced Salt Solution, non-essential amino acids, 2 mM L-glutamine, 1 mM sodium pyruvate, and 1500 mg/L sodium bicarbonate, 10% (v/v) FBS and antibiotics (50 U penicillin, 50μg streptomycin) in 5% CO2 at 37°C. After 36 hours, cells were synchronized at metaphase in the presence of 0.03μg of colcemid (Sigma) for 12 hours, and collected for staining at 48 hours post MAO treatment. Cells were washed in PBS, followed by a single 20 min. fixation and permeabilization step using 4% paraformaldehyde (Alfa Aesar, Ward Hill, MA) with 1% Triton-X (Calbiochem) in PBS. Additional PBS washes were completed prior to blocking with 7% donkey serum (Jackson ImmunoResearch Laboratories, Inc., West Grove, PA) in PBS for either 1 hour at RT or overnight at 4°C. A primary antibody against BUB1B (ab28193, Abcam, Cambridge, MA) was diluted 1:1000 in PBS with 1% donkey serum and cells were incubated overnight at 4°C. BUB1B antibody binding was detected using a 568-conjugated donkey, anti-rabbit Alexa Fluor secondary antibody (Thermo Fisher) at a 1:250 dilution with 1% donkey serum in PBS at RT for 1 hour in the dark. Cells were washed in PBS and the DNA stained with 1 μg/ml DAPI for 15 min. The coverslips with adherent cells were then mounted on slides using Prolong Diamond mounting medium (Invitrogen, Carlsbad, CA, USA). Immunofluorescence was visualized on a Nikon Eclipse Ti-U fluorescent microscope system and representative fluorescent images captured with a Nikon DS-Ri2 color camera. Using FIJI, background fluorescence was subtracted from the red (BUB1B) channel, followed by individual channels of each color for combination into a composite image. BUB1B immunostaining was visually assessed at each MAO concentration for 100 metaphase cells per treatment group.

### BUB1B knockdown and validation

Zygotes underwent cytoplasmic injection with 3’-carboxyfluorescein-labeled MAO at 20 hours post fertilization as described above. A concentration of 0.3 mM MAO was used based on previous findings that Std Control MAO at this concentration was the maximum which allowed normal blastocyst formation rates. Following microinjection, embryos were cultured up to the blastocyst stage as described above with or without imaging on the Eeva™ darkfield 2.2.1 microscope system. Upon developmental arrest, embryos were collected for immunostaining, gene expression analysis, or disassembled into single cells (as described above) for downstream analysis. To further validate MAO specificity, bovine embryos were co-injected with BUB1B modified mRNA at a concentration of approximately 3nl (75pg) of mRNA per embryo in addition to BUB1B MAO #1. The BUB1B coding sequence (CDS) was amplified from the plasmid, pcDNA5-EGFP-AID-BubR1 (Addgene #47330), followed by mutation of the MAO binding site using the Q5 site directed mutagenesis kit (NEB) according to the manufacturer’s instructions. Briefly, custom primers (forward: 5’-aaaaaagagggaGGTGCTCTGAGTGAAGCC-3’, and reverse: 5’-aactgcagccatATGGGATCCAGCTCTGCT-3’) were designed to mutate the region of the BUB1B CDS targeted by the MAO without affecting the amino acid sequence. Exponential amplification of the template plasmid using high fidelity DNA polymerase was followed by a single step phosphorylation, ligation and DpnI restriction enzyme digestion. NEB 5-apha competent cells were transformed with the mutated plasmid, followed by DNA miniprep isolation using QIAprep spin columns (Qiagen). Mutated plasmids were identified by Sanger sequencing performed by the ONPRC Molecular and Cellular Biology Core using a custom designed primer (TTGGTGAATAGCTGGGACTATG). Following identification and isolation, the mutated plasmid served as a template to synthesize a PCR product containing a T7 promoter using Platinum Taq (Invitrogen). Custom primers (forward: CTAGCTTAATACGACTCACTATAGGGAGCGCCACCATGGCTGCAGTTAAAAAAGAG, reverse: CAATCTGTGAGACTTGATTGCCTAGCTCACTGAAAGAGCAAAGCCCCAG) were designed for use with the T7 mMessage mMachine Ultra Kit as described above.

### Quantitative RT-PCR analysis

Gene expression was analyzed in non-injected, Std control MAO, or BUB1B MAO injected embryos using the BioMark Dynamic Array microfluidic system (Fluidigm Corp., So. San Francisco, CA, USA). All embryos were collected within 36 hours post fertilization as described above. Individual embryos were pre-amplified according to the manufacturer’s “two-step single cell gene expression” protocol (Fluidigm Corp.) using SuperScript VILO cDNA synthesis kit (Invitrogen), TaqMan PreAmp Master Mix (Applied Biosystems, Foster City, CA, USA), and gene-specific primers designed to span exons using Primer-BLAST (NCBI). Bovine fibroblasts and no RT template samples were used as controls. Pre-amplified cDNA was loaded into the sample inlets of a 96 × 96 dynamic array (DA; Fluidigm Corp.) and assayed in triplicate. A total of 10 reference genes were assayed for use as relative expression controls. Cycle threshold (Ct) values were normalized to the two most stable housekeeping genes (*RPL15* and *GUSB*) using qBase^+^ 3.2 software (Biogazelle; Ghent, Belgium). Calculated normalized relative quantity (CNRQ) values were averaged across triplicates + the standard error and graphed using Morpheus (https://software.broadinstitute.org/morpheus/).

### Statistical analysis

To determine statistical differences between MAO concentrations in MDBK cells, log-binomial modeling using the Generalized Estimating Equations approach was performed and Tukey adjusted p-values reported to adjust for multiple comparisons. From the qRT-PCR results, averaged CNRQ values of each gene was compared across embryo groups using the Mann-Whitney U-test. An unadjusted p-value<0.05 was considered statistically significant.

## ACKNOWLEDGEMENTS

We gratefully acknowledge Dr. Tom Spencer at the University of Missouri-Columbia for the Madin-Darby Bovine Kidney (MDBK) epithelial cells. K.E.B. was supported by the NIH/NICHD Postdoctoral Individual National Research Service Award (5F32HD095550-01). B.L.D. was supported by the P.E.O. Scholar Award, N.L. Tartar Research Fellowship, and T32 Reproductive Biology NIH Training Grant (T32 HD007133). The authors acknowledge the support of the Oregon National Primate Research Center (ONPRC) Integrated Pathology Core for confocal microscopy (supported by S10RR024585) that operates under the auspices of the ONPRC NIH/OD core grant (P51OD011092). This work was supported by OHSU/ONPRC start-up funds (to SLC) and the NIH/NICHD (R01HD086073-A1). The content of this paper is solely the responsibility of the authors and does not necessarily represent the official views of the NIH.

## AUTHOR CONTRIBUTIONS

Conceptualization, K.E.B. and S.L.C; Methodology, K.E.B., B.L.D., and S.L.C.; Software, K.E.B., B.D., and M.Y.Y.; Validation, K.E.B. and S.L.C.; Formal Analysis, K.E.B. and S.L.C.; Investigation, K.E.B. and B.L.D.; Data Curation, S.S.F. and S.L.C.; Writing – Original Draft, K.E.B. and S.L.C.; Writing – Review & Editing, K.E.B., B.L.D., B.D., M.Y.Y., S.S.F., L.C., and S.L.C; Visualization, K.E.B., B.D., and M.Y.Y.; Supervision, S.S.F., L.C., and S.L.C.; Project Administration, S.L.C.; Funding Acquisition, K.E.B., L.C., and S.L.C.

## Supplemental Materials

### Supplemental Figures

**Supplemental Figure S1.**
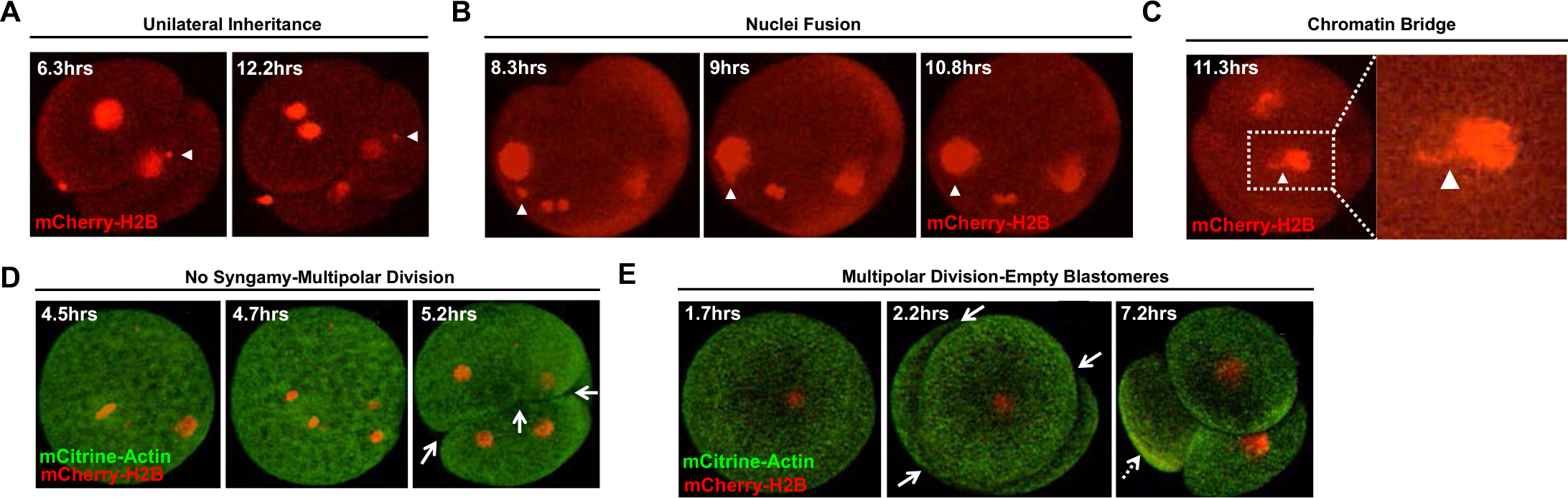
Additional live-cell images representative of embryos with different phenotypes. Live-cell confocal microscopy of bovine zygotes microinjected with fluorescently labeled modified mRNAs to visualize DNA (Histone H2B-mCherry; red) and distinguish blastomeres (Actin-mCitrine; green) during the first three mitotic divisions. **(A)** Examples of other embryos with micronuclei that undergo unilateral inheritance, **(B)** fuse back with the primary nucleus, or **(C)** form a chromatin bridge (white arrowheads). **(D)** Images of additional embryos that bypassed pronuclear fusion (syngamy) prior to a multipolar division (white solid arrows) to produce blastomeres with uniparental origins and/or **(E)** no apparent nuclear structure (white dashed arrows). Individual frames are represented in hours (hrs) from the start of imaging.

**Supplemental Figure S2.**
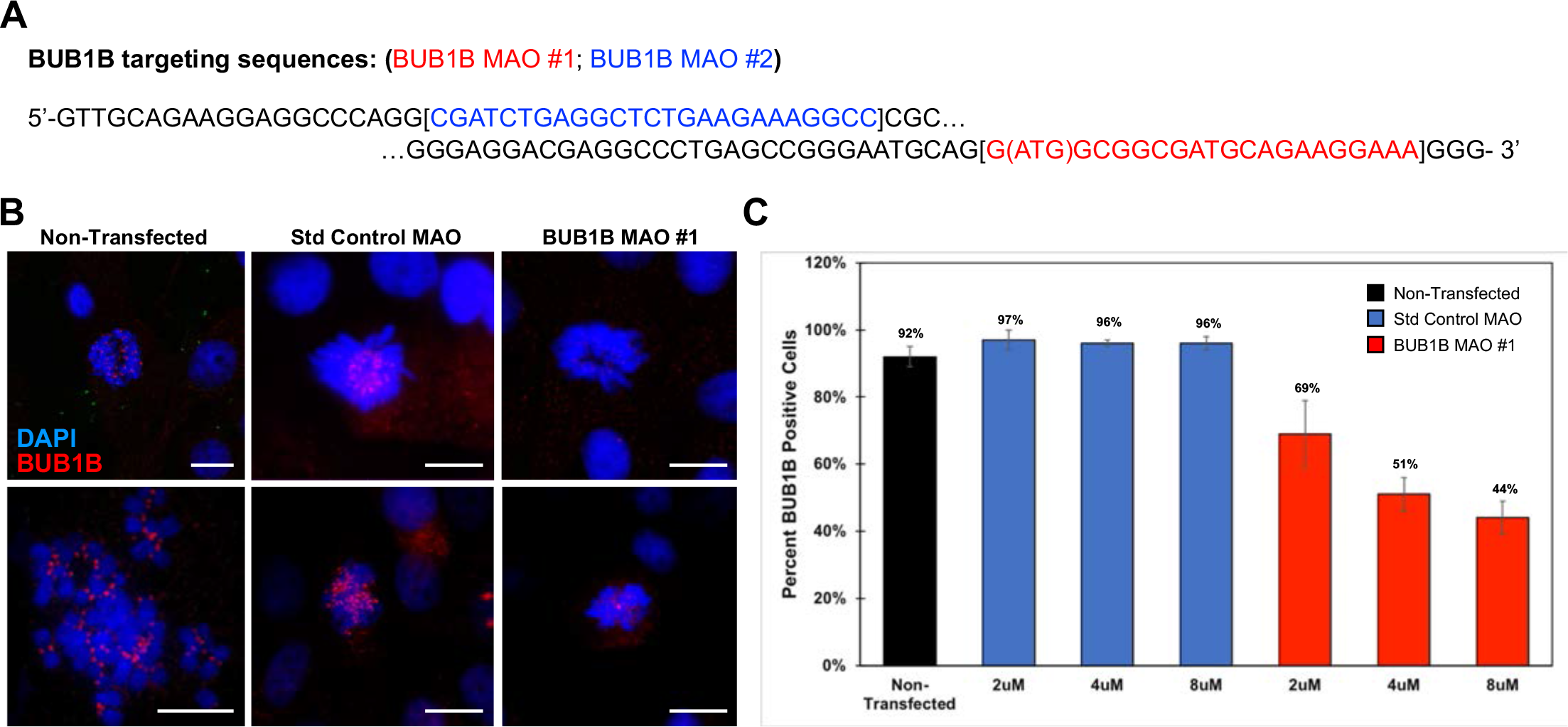
BUB1B MAO design and knockdown efficiency. **(A)** DNA sequences of two non-overlapping MAOs designed to target the ATG start site (shown in red, BUB1B MAO #1) and the 5’ UTR (depicted in blue, BUB1B MAO #2) of BUB1B. **(B)** BUB1B knockdown efficiency was assessed in synchronized MDBK cells following 48 hours of treatment with 3μl/ml of colcemid alone (non-transfected), the Std control MAO, or BUB1B MAO #1 via immunofluorescence. BUB1B protein expression was analyzed in DAPI stained (blue) MDBK cells. Note the lack of or reduced number of BUB1B positive foci (red) in the BUB1B MAO #1 treated cells compared to the controls; Scale bar = 20μm. **(C)** Bar graph showing the percentage of MDBK cells in metaphase with BUB1B expression after colcemid treatment (black) or transfection with different concentrations (2, 4, and 8 μM) of the Std control MAO (blue) or BUB1B MAO #1 (red). While the number of cells exhibiting BUB1B positive foci was similar between the non-transfected and Std MAO controls, a dose-dependent decrease in BUB1B expression was observed following BUB1B MAO #1 treatment.

**Supplemental Figure S3.**
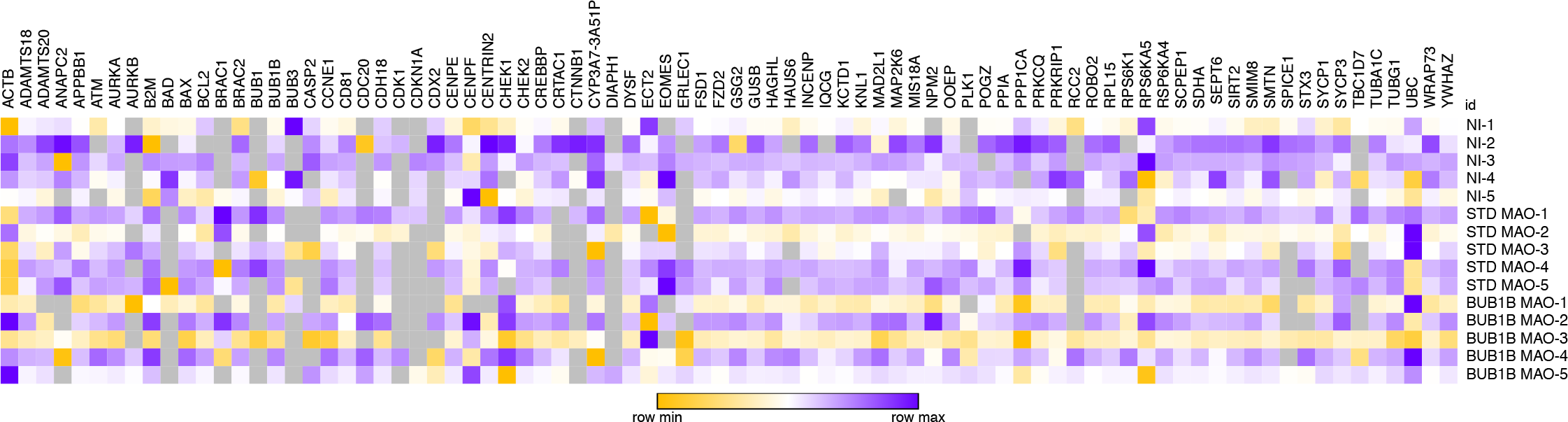
Comprehensive assessment of gene expression patterns in zygotes. Heat map of all mitotic, cell cycle, developmentally-regulated, and cell survival genes assessed in individual BUB1B MAO #1 versus non-injected and Std Control-injected MAO bovine zygotes via microfluidic qRT-PCR. Cycle threshold (Ct) values were normalized to the most stable reference genes (RPL15 and GUSB) across embryo groups and presented as the average. Gray squares indicated no expression, whereas yellow, white, and purple squares correspond to low, medium, and high expression, respectively.

### Supplementary Tables

**<u>Supplemental Table S1.</u> Sequencing statistics of all embryonic and control samples.** A table depicting the number or percentage of reads following de-multiplexing of embryonic (with embryo stage) and fibroblast samples at each step of the post-sequencing process, including adaptor removal, repeat masking, genome mapping, and quality assessment. The sequencing kit used and whether single- or paired-end is also included.

**Supplemental Table S3.**
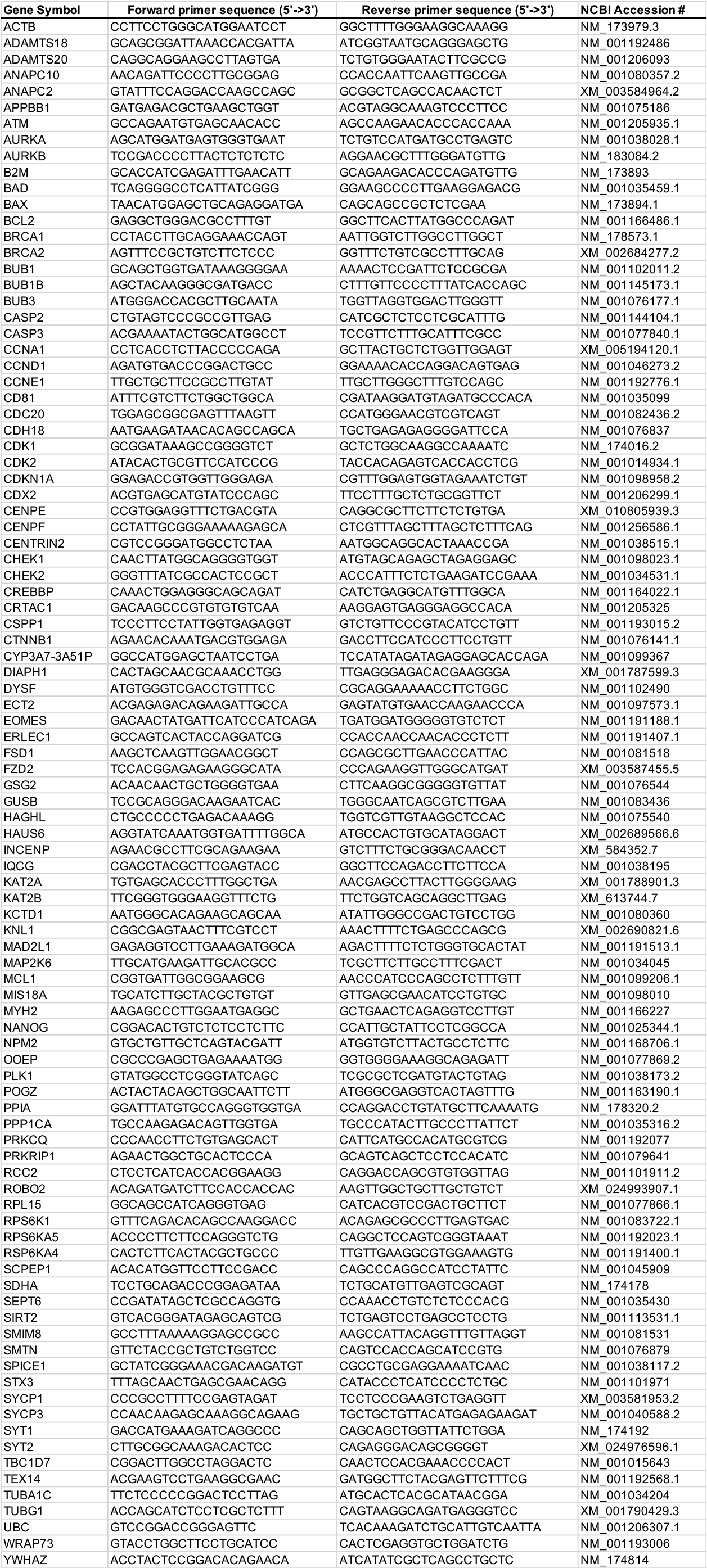
List of all genes with primers analyzed by qRT-PCR in zygotes. A table of the genes analyzed by microfluidic qRT-PCR in non-injected bovine zygotes and following Std Control MAO versus BUB1B MAO #1 microinjection. Included is the sequence of the forward and reverse primer used for amplification as well as the NCBI accession number of each gene.

### Supplemental Movies

**<u>Movie S1</u>. Live-cell fluorescent imaging of early cleavage divisions.** Bovine zygotes were microinjected with fluorescently labeled modified mRNAs to mCitrine-Actin (green) and mCherry-Histone H2B (red) to distinguish blastomeres and DNA, respectively, and early mitotic divisions visualized by live-cell confocal microscopy. Note the micro-/multi-nuclei in embryos #3, #4, and #11, chromatin bridge in embryo #1, lack of syngamy in embryos #3 and #11, multipolar divisions in embryos #1, #3-6, #11, and #15, and production of empty blastomeres in embryos #5 and #15.

**<u>Movie S2</u>. MCC-deficient embryos struggle to divide.** A bovine zygote following BUB1B MAO microinjection attempted to divide by forming multiple cleavage furrows, but never successfully completed cytokinesis.

**<u>Movie S3</u>. Multipolar divisions are observed in MCC-deficient embryos.** Certain bovine zygotes were able to undergo cytokinesis even with BUB1B knockdown, but these divisions were abnormal with multipolar cleavage.

**<u>Movie S4</u>. MCC deficiency causes blastomere asymmetry.** Besides abnormal divisions, BUB1B-injected bovine embryos often exhibited blastomere asymmetry following the multipolar cleavage.

